# Accurate determination of breed origin of alleles in a simulated smallholder crossbred dairy cattle population

**DOI:** 10.1101/2024.04.12.589204

**Authors:** Berihu Welderufael, Isidore Houaga, Chris R Gaynor, Gregor Gorjanc, John M Hickey

## Abstract

**Background:** Accurate assignment of breed origin of alleles at a heterozygote locus may help to introduce a resilient or adaptive haplotype in crossbreeding. In this study, we developed and tested a method to assign breed of origin for individual alleles in crossbred dairy cattle. After generations of mating within and between local breeds as well as the importation of exotic bulls, five rounds of selected crossbred cows were simulated to mimic a dairy breeding programme in the low- and middle-income countries (LMICs). In each round of selection, the alleles of those crossbred animals were phased and assigned to their breed of origin (being either local or exotic).

**Results:** Across all core lengths and modes of phasing (with offset or no), the average percentage of alleles correctly assigned a breed origin was 95.76%, with only 1.39% incorrectly assigned and 2.85% missing or unassigned. On consensus, the average percentage of alleles correctly assigned a breed origin was 93.21%, with only 0.46% incorrectly assigned and 6.33% missing or unassigned. This high proportion of alleles correctly assigned a breed origin resulted in a high core-based mean accuracy of 0.99 and a very high consensus-based mean accuracy of 1.00. The algorithm’s assignment yield and accuracy were affected by the choice of threshold levels for the best match of assignments. The threshold level had the opposite effect on assignment yield and assignment accuracy. A less stringent threshold generated higher assignment yields and lower assignment accuracy.

**Conclusions:** We developed an algorithm that accurately assigns a breed origin to alleles of crossbred animals designed to represent breeding programmes in the LMICs. The developed algorithm is straightforward in its application and does not require prior knowledge of pedigree, which makes it more relevant and applicable in LMICs breeding programmes.

## Background

Dairy cattle production in low- and middle-income countries (LMICs) is characterised by low-input and low-output production systems. To increase the milk productivity of dairy cattle, crossbreeding between the high-producing breeds of developed countries and the low-producing, but resilient breeds of LMICs has been practised for decades. Crossbreeding, either via the importation of semen from elite bulls or the use of imported bulls, has substantially increased milk production and farmers’ income [1]. However, this genetic gain has not always been observed, and overreliance on import without judicious use of best alleles is not expected to deliver the best possible genetic gains.

In many LMICs, including those in Eastern Africa, efforts are being undertaken to establish sustainable breeding programmes for long-term genetic gains with a focus on smallholder farmers [2]. In partnership with government and non-government organizations, projects like the African Dairy Genetic Gains (ADGG, https://africadgg.wordpress.com) have been able to import and provide improved dairy genetics to smallholder farmers in the Eastern Africa. However, because of the differences in environmental factors and breeding infrastructure, the importation and provision of improved genetics have not yet been sustainable and successful [2]. Instead, such crossbreeding practices have led to haphazardly admixed cattle populations with no or poor pedigree records [2].

For a sustainable breed improvement through genetic intervention and for the appropriate design of breeding programmes, accurate breed identification, on both the level of the individual and of the individual genetic variant, is important. In livestock populations with little or no pedigree records, the use of genomic data could be transformational in determining breed composition and establishment of breeding programmes [2]. Estimates of breed composition and the breed origin of alleles from genomic data is superior to estimates from pedigree data due to invariably missing or inaccurate records and deviations from expected compositions due to Mendelian sampling [3,4]. Especially in populations with complex ancestries like the dairy cattle in the Eastern Africa, genomic data and knowledge of breed composition is essential to evaluate the performance and adaptability of the crossbreds [4], and to predict the effectiveness of any foreign germplasm in the production systems.

Selection, genetic discovery and management decisions can be aided by determining the breed origin of alleles, particularly for genetic variants that only occur in one of the constituent populations of crossbred animals [5]. Unlike determining the average breed composition of an individual, determining the breed origin of an individual’s haplotypes and associated alleles can allow breed-specific genetic evaluations to be conducted, which can increase the accuracy of genetic selection, particularly when the linkage disequilibrium pattern is different in the two breeds [6]. Thus, recent studies in admixed cattle populations have shown that the Breed Origin of Allele (BOA) method has increased the accuracy of genomic prediction [7,8].

Using only genomic data and no pedigree data, Vandenplas et al. [5] developed an approach that traces haplotypes of crossbred animals and assigns each allele of the haplotypes to their breed of origin. To develop the algorithm that assigns alleles of crossbreds a breed origin, they simulated a three-way pig-crossbreeding programme with five generations of random selection. They evaluated the accuracy of the algorithm and reported that more than 90% of alleles of crossbred animals were correctly assigned a breed origin. Thus, for up to 10% of all alleles of crossbred animals, they could not assign a breed origin. However, accurate determination of the breed origin of alleles of crossbred populations is very important to estimate breed- specific effects of alleles when performing genomic evaluations [9]. If we could accurately assign breed origin for alleles at heterozygote loci of crossbred animals, we may be able to detect which haplotypes should be promoted to genetically improve dairy cows in the LMICs.

In the current study, we developed an algorithm to assign a breed of origin for alleles in crossbred dairy cattle and tested it on a simulated smallholder dairy cattle population dataset. To resolve the breed origin of alleles, the algorithm aligns the haplotypes of crossbred dairy cows to the haplotypes of likely constituent breeds, i.e., imported (exotic) and/or local breeds and assigns the breed of origin based on the best match. We then evaluated the algorithm’s accuracy using a simulated crossbreeding programme designed to mimic the ADGG smallholder genotype data. The average percentage of alleles correctly assigned a breed origin was 95.76%, resulting in a high core-based mean accuracy of 0.99 and a very high consensus-based mean accuracy of 1.00. The developed algorithm does not require prior pedigree knowledge and is, hence, straightforward to apply in LMIC breeding programmes.

## Methods

The design of the breeding programme and development of the allele assignment algorithm involved two steps.

1. We designed a breeding programme and simulated genotype data on which we tested the algorithm’s performance. The simulated genotype data had an ancient cattle founder that is assumed to have split into African (local) and European (exotic) cattle populations. After generations of mating within and between local breeds and the importation of exotic bulls, crossbred dairy cows were created to mimic the dairy cows kept by smallholders in the LMICs.
2. We developed an allele assignment algorithm that traces haplotypes and assigns a breed origin for each allele of the crossbred cows. The haplotypes are phased and defined for different core lengths to improve the accuracy and efficiency of the allele assignment algorithm.

The following subsections describe the details for simulating and phasing genotypes and developing the allele assignment algorithm.

### Simulation of genotype data

Genotype and haplotype data for an ancient cattle breed were simulated using the AlphaSimR package [10], designed for stochastic simulations of breeding programmes. A total of 2500 individual animals with a genome structure of 1000 SNPs in one autosomal chromosome were simulated. The ancient cattle breed split into two, each representing an exotic breed and an indigenous breed. The indigenous breed further split into four more closely related local founder populations. Variation in the demographic history of these founder populations were accounted for in the simulated biallelic haplotypes of the breeds using the Markovian Coalescent Simulator (MaCS) software [11] embedded in the AlphaSimR package [10]– [See

Additional file 1, Script S1] for details. As described in the AlphaSimR, the genotypes and haplotypes of the descendants, i.e., the crossbred animals, were then derived from these haplotypes using simulated mating between the exotic and local breeds. After within and between breed random mating of indigenous animals for 10 generations, the 1000 best females were selected on genetic merit of a single hypothetical trait with a small amount of dominance (mean dominance degree of 0.1 and variance of dominance degree of 0.1) and heritability of h^2^=0.3. The 1000 selected local cows were then mated to 25 imported Holstein bulls to produce the first crossbred animals (crossbred1). The local cows were allowed to calve twice producing a total of 2000 offsprings with the assumption of 1000 female and 1000 male calves. The breeding programme continued by using all the 1000 female calves (crossbred1) as replacement heifers and mating these to 25 newly imported Holstein bulls to produce the next crossbred cows (crossbred2), while both exotic and local populations were kept as purebred and source of purebred animals. This importation of exotic bulls and mating to the crossbred cows was repeated for up to five rounds of selections, hereafter referred as generations (Fig. 1). Simulated genotype and haplotype data were generated in 10 replicates.

**Figure 1.**
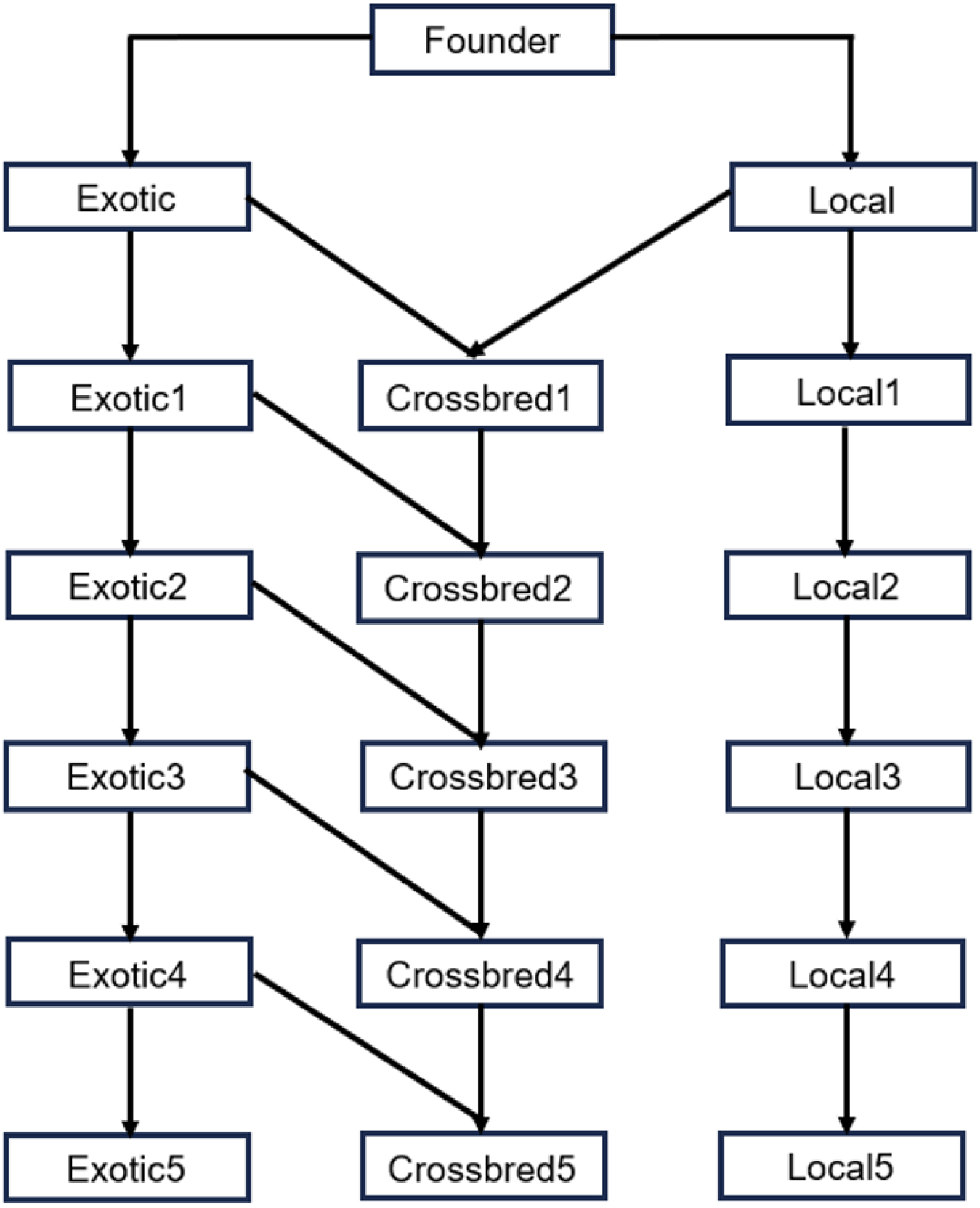
Schematic representation of the simulated breeding programme. A founder population on the top of the figure is split into exotic and local breeds.

### Genetic structure of the simulated SNP genotype data

To assess the genetic similarity between the founders and developed crossbred animals, we performed principal component analysis (PCA) of SNP genotypes on the simulated data. The PCA was performed using the prcomp command of the R statistical software [12].

### Phasing of simulated genotype data

True simulated genotype and haplotype data enabled us to calculate the phasing yield and allele assignment accuracy. From the genotype data, haplotypes were reconstructed and compared with the simulated haplotypes. The reconstruction of possible haplotypes from the genotype data via phasing was performed using the software AlphaPhase [13]. Different core and tail lengths govern the length of desired haplotype segments used to phase the alleles in the genotype data. As illustrated in Fig. 2, a core is a string of consecutive SNP loci used to phase a given genome region [13].

**Figure 2.**
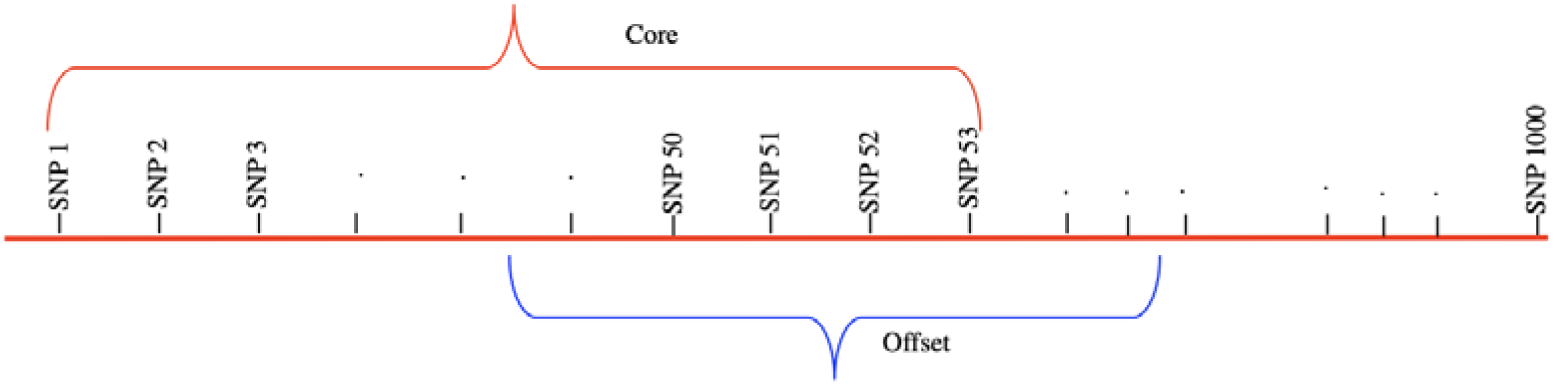
Illustration of a core and offset. Phasing was performed in two modes: either using the whole length of a core or by moving it forward 50% of the core length (offset) to define the begging of a given core.

Phasing of the simulated genotype data was performed using a wide range of core and tail lengths. Preliminarily analyses suggested that core lengths of 100 to 280 SNPs would yield optimum allele assignments. Therefore, for the final analyses, we defined 10 different core lengths centred around 280 SNPs (Table 1) and phasing was performed for each core length both in the offset and no-offset modes of the AlphaPhase [13]. We moved 50% of the core length forward to define Offset. In total, there were 2000 scenarios: 10 (replicates) x 10 (core lengths) x 10 (thresholds) x 2 (offset or no offset modes).

**Table 1.**
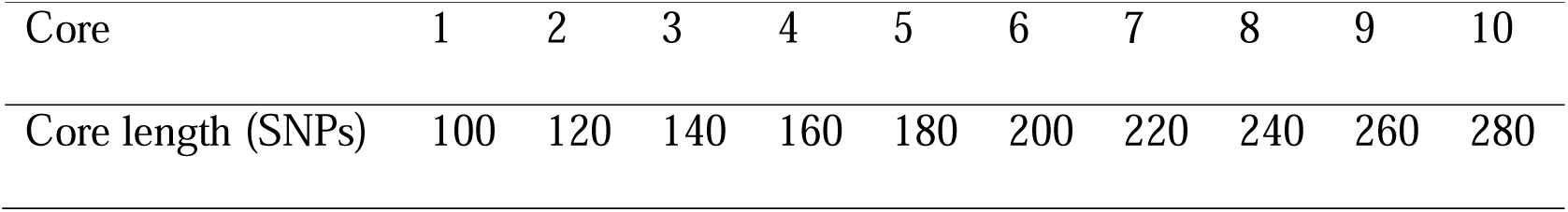
Core lengths (in terms of numbers of SNPs) used to phase the genotype data.

|  |  |  |  |  |  |  |  |  |  |  |
| --- | --- | --- | --- | --- | --- | --- | --- | --- | --- | --- |
| Core | 1 | 2 | 3 | 4 | 5 | 6 | 7 | 8 | 9 | 10 |
| Core length (SNPs) | 100 | 120 | 140 | 160 | 180 | 200 | 220 | 240 | 260 | 280 |

### Development of allele assignment algorithm

To develop the allele assignment algorithm, we defined 10 different core lengths (Table 1). The alleles of crossbred animals were assigned a breed origin for each core length, and we call this core-based allele assignment. In the core-based assignment each allele could be assigned a breed origin as many as the different core lengths defined. If breed origin assignments of an allele were not the same across the different cores the most frequent breed assignment was considered as a consensus breed origin of an allele.

### Core-based allele assignment

Haplotype libraries were simulated based on the phased purebred individuals in each population. The assignment algorithm takes phased genotypes for individuals in the crossbred population as inputs, along with haplotype libraries for the indigenous and exotic populations (Fig. 3). To perform allele assignment, we determined whether the exotic or local haplotype contained the best matching haplotype, i.e. the haplotype with the fewest number of markers than the target haplotype. The haplotype is then assigned as originating from that haplotype library. If both haplotype libraries contain an equally good match, then the haplotype is set to missing. For example, in Fig. 3, the haplotype with a core length of 10 SNPs of the individual animal should be assigned to the local haplotype as it displays the least error matches with the last core in the local haplotype library.

**Figure 3.**
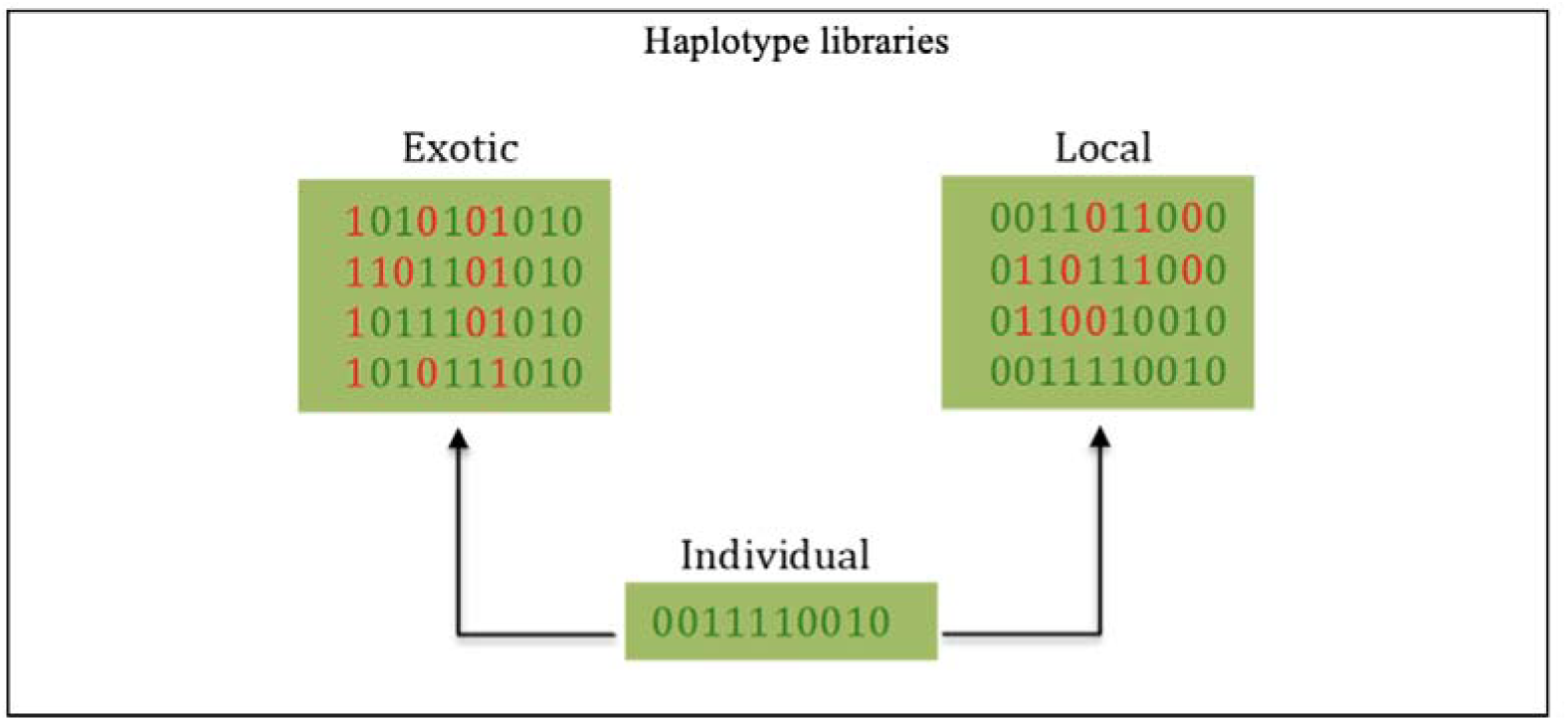
Haplotype libraries based on a core length of ten SNPs. To assign origin to the haplotype of an individual (bottom genotype sequence), the algorithm searches for the best match in each position in the exotic (top left genotype sequence) and local (top right genotype sequence) haplotypes. In this case, the individual’s haplotype should be assigned as a local haplotype because the local haplotype library contains the haplotype with the fewest number of errors, i.e., mismatches (red).

### Consensus allele assignment

Allele assignment was compared in each phased genotype and each scenario. Phasing of simulated genotype data was performed in two modes: either using the whole length of a core or by moving it forward 50% of the core length (offset) to define the beginning of a given core (see next section). Assignment was performed across multiple core lengths and two modes of phasing (no offset and offset). Assignment results of each core and mode of phasing were compared and merged across cores to calculate consensus-based assignment. Merging was done by taking a consensus estimate of the breed of origin across multiple cores. The most frequently observed assignment across all the replicates, core lengths, and phasing modes was then taken as the consensus-based assignment.

To optimise and fine-tune the algorithm’s sensitivity, we applied 10 different thresholds for best SNP count of match of haplotypes (Table 2). When the threshold was 0.9, this meant that the breed assignment for the allele needed to be consistent across 90% of the cores, otherwise the assignment was set to missing. To elaborate a threshold of 50%, an allele would have been assigned a breed origin of “A” if the allele had been assigned to breed “A” in more than 50% times of the assignments across all the different core lengths and phasing modes. In every generation, every allele of the crossbred animals was assigned a breed origin in at least 2000 scenarios and results were merged to calculate consensus assignment.

**Table 2.** The different thresholds used for the best count of match of haplotypes.

| Threshold | 1 | 2 | 3 | 4 | 5 | 6 | 7 | 8 | 9 | 10 |
| --- | --- | --- | --- | --- | --- | --- | --- | --- | --- | --- |
| %Matched | 0.50 | 0.55 | 0.60 | 0.65 | 0.70 | 0.75 | 0.80 | 0.85 | 0.90 | 0.95 |

### Performance of the allele assignment algorithm

To evaluate the performance of the allele assignment algorithm, assignment yield and assignment accuracy were assessed in the following ways:

1. %Correct: the percentage of correctly assigned alleles was computed by comparing the algorithm-derived breed origin with the true breed origin of alleles traced with the “pullIbdHaplo()” function of the AlphaSimR [10].
2. %Incorrect: the percentage of alleles across all scenarios that were incorrectly assigned and was computed by comparing the algorithm-derived breed origin with the true breed origin of alleles traced with the “pullIbdHaplo()” function of the AlphaSimR [10].
3. %Unassigned: the percentage of alleles that were not assigned, including missing or unknown breed origin; and
4. Accuracy: the accuracy of assigned alleles, calculated as the ratio of correctly and incorrectly assigned alleles. We used the proportion of correctly assigned alleles as an allele assignment accuracy metric for each core and tail lengths.

## Results

### Genetic structure of the simulated SNP genotype data

Principal component analysis (PCA) of the simulated SNP genotype data separated the crossbreds from the founder breeds (local and exotic breeds). As shown in the PCA plot (Fig. 4a), the first generation of crossbred animals (crossbred1) were positioned in between the founder populations (exotic and local). The PCA plot further revealed the genetic sub-structure from the crossbreeding programme. As we continued the crossbreeding and increased the proportion of exotic genotypes, the crossbreds and the exotic breed were observed to converge into a single cluster (Fig. 4b).

**Figure 4.**
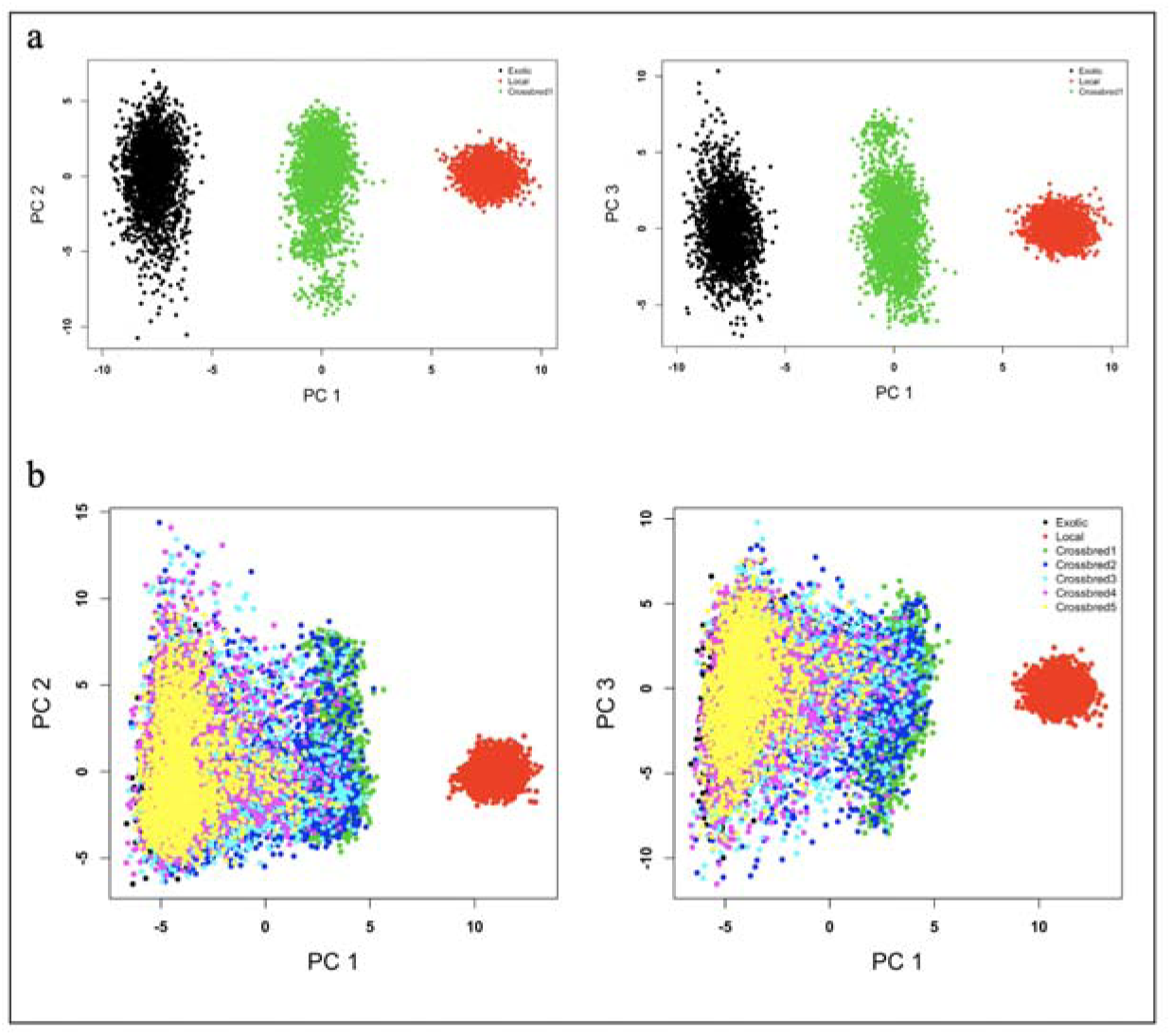
Plot of principal component analysis of SNP genotypes (PC1 vs. PC2 and PC1 vs. PC3). Showing the genetic data structure of the founders and the first crossbred cows (a) and of all animals across generations (b).

### Allele assignment yield and accuracy

#### Allele assignment for each core

The average number of alleles in the crossbred animals assigned a breed origin is given in Table 3. The highest average number of unassigned alleles (29 out of 1000 SNPs) was observed in the first generation of the crossbred animals (crossbred1). The number of unassigned alleles decreased as the crossbreeding continued and the distance between the local founders and the crossbreds decreased. For example, in crossbred5, where the germplasm is upgraded to almost the exotic breed, 23 out of 1000 SNPs were unassigned (Table 3).

**Table 3.** Assignment yield and average number of alleles in crossbred cows assigned to local, exotic or not assigned at all.

| Crossbred | Local | Exotic | Unassigned | Assignment<br>Yield |
| --- | --- | --- | --- | --- |
| Crossbred1 | 486 | 486 | 29 | 0.95 |
| Crossbred2 | 246 | 730 | 24 | 0.95 |
| Crossbred3 | 123 | 853 | 24 | 0.96 |
| Crossbred4 | 61 | 916 | 24 | 0.96 |
| Crossbred5 | 29 | 947 | 23 | 0.97 |
| Mean | 189 | 786 | 25 | 0.96 |

The genetic distance and core lengths had a clear effect on the phasing and assignment yield. For longer core lengths (core length of 220-280 SNPs), we observed a more concise and higher phasing yield (Fig. 5a). A core length of 200 SNPs was observed to be optimal for allele assignment yield (Fig. 5b). The overall average allele assignment accuracy was 0.99 (Table 4). On average, more than 95% of the assigned alleles in the crossbred animals were correctly assigned, with only less than 2% of incorrectly assigned alleles (Table 4). Both, the incorrectly assigned and unassigned proportion of alleles, either because of missing or ambiguity, were less than 5% (Table 4).

**Figure 5.**
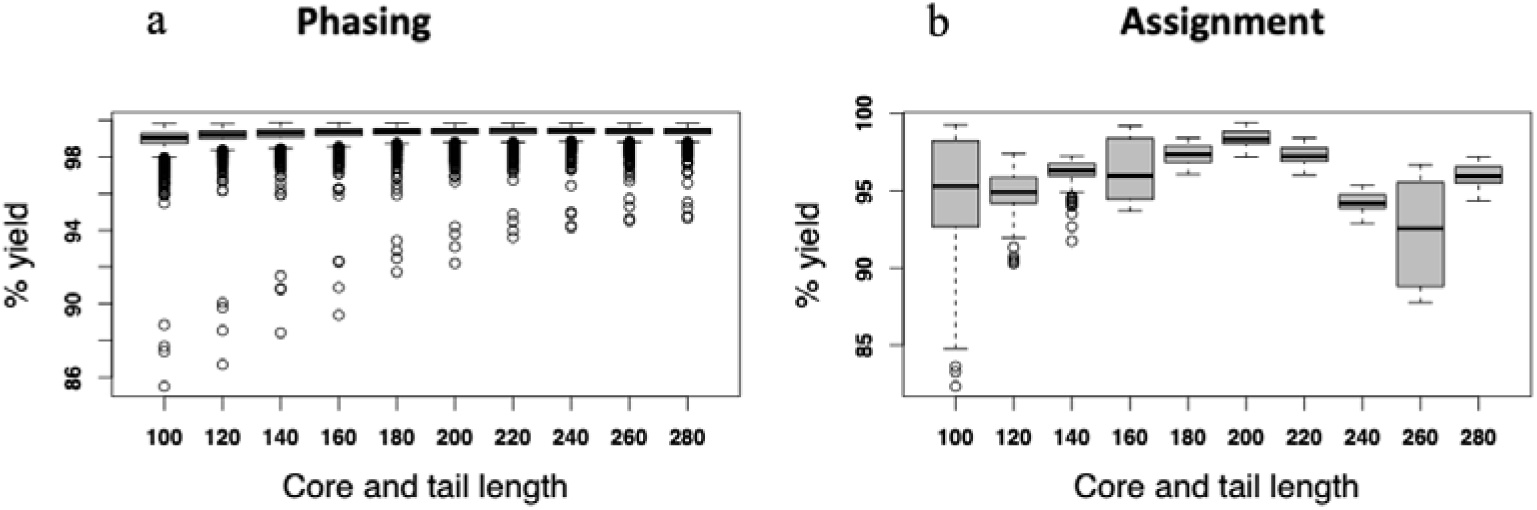
Effect of core length on assignment yield. Phasing yield (a) was very high for all core lengths but more concise for longer core lengths (core length of 220-280 SNPs). The assignment yield (b) was optimal for a core length of 200 SNPs.

**Table 4.** Percentages of alleles correctly assigned a breed origin (%Correct), incorrectly assigned (%Incorrect), missing or unassigned (%Unassigned), and accuracy of assignment (Accuracy) for each core-length (Core)

| Core | %Correct | %Incorrect | %Unassigned | Accuracy |
| --- | --- | --- | --- | --- |
| 100 | 94.70 | 1.35 | 3.95 | 0.99 |
| 120 | 94.89 | 1.12 | 3.99 | 0.99 |
| 140 | 96.14 | 1.11 | 2.75 | 0.99 |
| 160 | 96.40 | 1.16 | 2.44 | 0.99 |
| 180 | 97.39 | 1.25 | 1.35 | 0.99 |
| 200 | 98.35 | 1.36 | 0.29 | 0.99 |
| 220 | 97.27 | 1.46 | 1.27 | 0.99 |
| 240 | 94.22 | 1.54 | 4.24 | 0.98 |
| 260 | 92.32 | 1.69 | 5.98 | 0.98 |
| 280 | 95.93 | 1.84 | 2.23 | 0.98 |
| Mean | 95.76 | 1.39 | 2.85 | 0.99 |

#### Consensus allele assignment across all cores

On consensus, the average percentage of incorrectly assigned alleles was nearly zero (Table 5). The overall mean consensus-based assignment accuracy (accuracy = 1, Table 5) was higher than the overall mean core-based assignment accuracy (accuracy = 0.99, Table 4).

**Table 5.** Consensus-based percentages of alleles correctly assigned (%Correct), incorrectly assigned (%Incorrect), missing or unassigned (%Unassigned) a breed origin, and accuracy of assignment (Accuracy) across all core-lengths and generation for each threshold.

| Threshold | %Correct | %Incorrect | %Unassigned | Accuracy |
| --- | --- | --- | --- | --- |
| 0.50 | 98.40 | 1.60 | 0.00 | 0.98 |
| 0.55 | 98.16 | 0.78 | 1.06 | 0.99 |
| 0.60 | 97.84 | 0.66 | 1.50 | 0.99 |
| 0.65 | 96.26 | 0.42 | 3.32 | 1.00 |
| 0.70 | 95.79 | 0.36 | 3.86 | 1.00 |
| 0.75 | 93.56 | 0.25 | 6.19 | 1.00 |
| 0.80 | 92.94 | 0.20 | 6.86 | 1.00 |
| 0.85 | 89.10 | 0.12 | 10.78 | 1.00 |
| 0.90 | 88.41 | 0.10 | 11.48 | 1.00 |
| 0.95 | 81.67 | 0.08 | 18.26 | 1.00 |
| Mean | 93.21 | 0.46 | 6.33 | 1.00 |

#### Effect of admixture level and thresholds on assignment yield and accuracy

The threshold level had the opposite effect on assignment yield and accuracy (Fig. 6). Increasing the threshold decreased the assignment yield and increased the accuracy, whereas a less stringent threshold generated higher assignment yields. Increasing the threshold stringency further reduced the assignment yield (Fig. 6a). On the contrary and as expected, the less stringent threshold reduced the accuracy (Fig. 6b).

**Figure 6.**
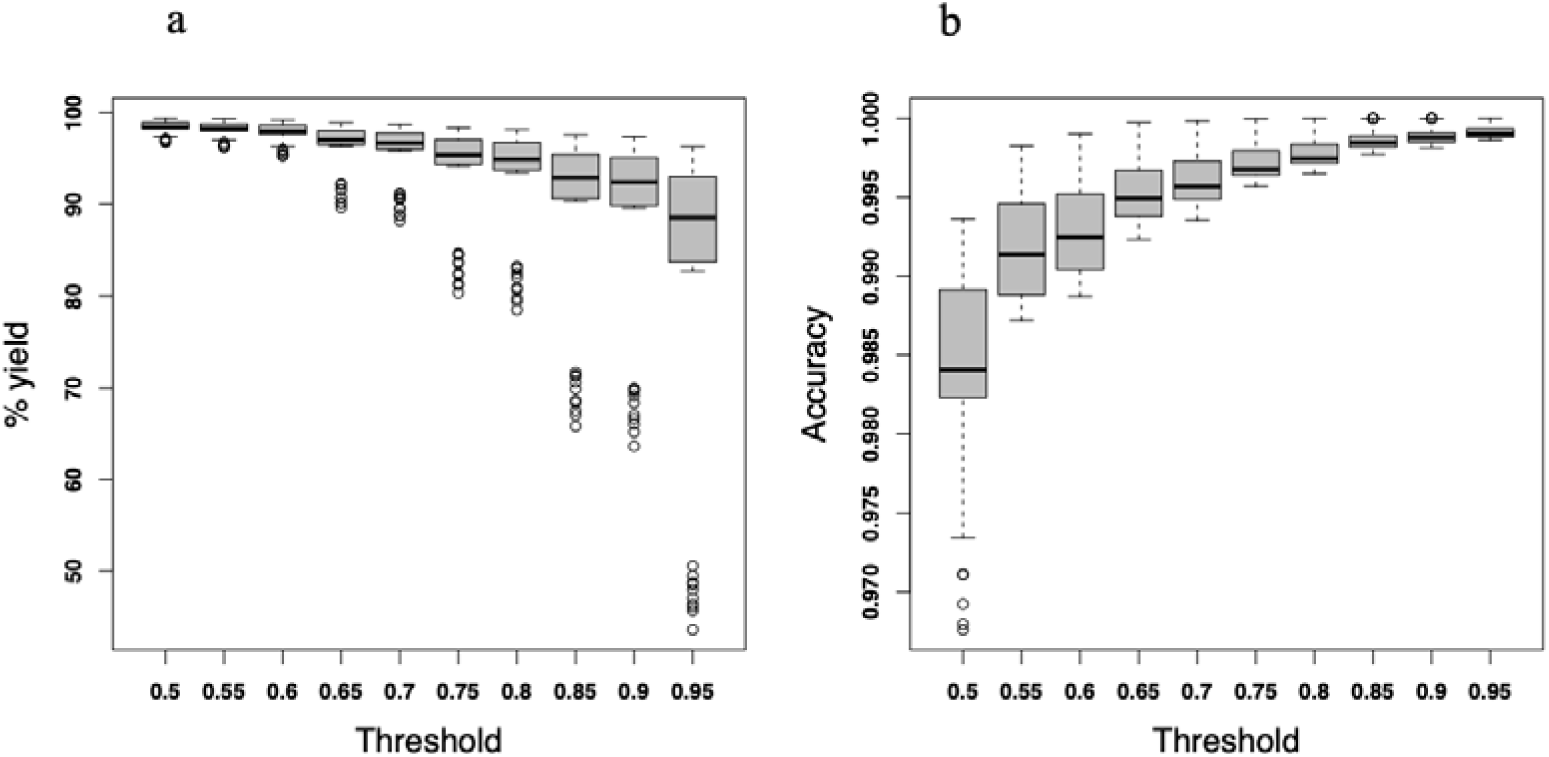
Percentage of allele assignment yield (a) and accuracy (b) of assignment. Using the consensus-based allele assignment algorithm as a function of threshold level

The effect of admixture level on assignment yield and accuracy was not as clear as that of threshold level. However, the assignment yield appeared to increase from the first to the later generations of crossbreds (Fig. 7a). On the other hand, the higher threshold stringency decreased the assignment yield (Fig. 7b).

**Figure 7.**
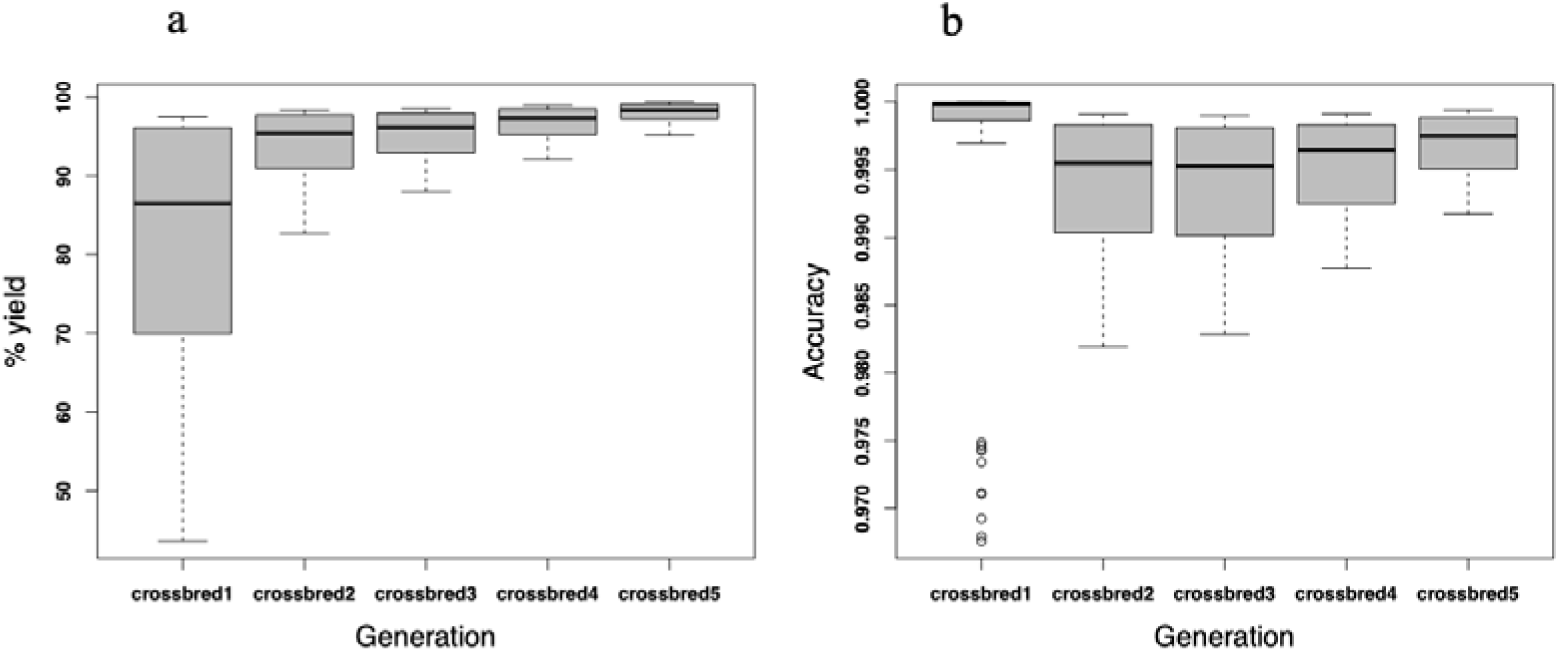
Percentage of allele assignment yield (a) and accuracy (b) of assignment. Using the consensus-based allele assignment algorithm as a function of crossbreeding (admixture) level.

## Discussion

In low- and middle-income countries (LMICs), such as those in Eastern Africa, a large proportion of dairy production is carried out by smallholders who keep fewer than 10 cattle [14]. These cattle are mostly crosses between indigenous African breeds and exotic dairy breeds, with little phenotypic or pedigree data available [14]. Despite the need and efforts to increase the productivity of those dairy cattle, it has not been possible to implement conventional breeding programmes in these populations. In populations with no or poor pedigree and phenotype records, genomic selection and other novel methods, such as an efficient algorithm to assign the breed origin of alleles in those crossbred animals, are of interest. To evaluate the performance and adaptability of the crossbreds in the LMICs, methods to accurately identify the breed origin of alleles on both the individual level and the individual genetic variant are important. Such methods could also provide ways to predict the effectiveness of foreign germplasm in a low-input production system [4]. For the smallholder farmers in Eastern Africa, providing methods to assign a breed origin of alleles would enable better choice of exotic bulls to introduce and which local bulls to use to sustainably harness the genetics of local adaptation traits of the indigenous breeds and the high milk yield potential of exotic dairy breeds.

Different genomic tools and algorithms [5,9] have been developed to assign a breed origin to alleles in crossbred pig populations without needing pedigree records. Using simulated genotype data, we have developed an algorithm to assign alleles a breed origin in a dairy cattle breeding programme that would represent haphazardly admixed local cows and imported exotic bulls as commonly practised in LMICs. As shown in Fig. 1, we used the exotic bulls as a source of purebred genotype data to cross with the admixed local cows for five subsequent generations. The simulated genotypes of exotic purebred and local admixed breeds were phased and the origins of haplotypes and associated alleles of the newly created crossbred cows were assigned a breed origin. In agreement with earlier studies in crossbred pig populations [5,9], our results demonstrated that alleles of admixed crossbred cattle populations could be accurately assigned a breed origin without the need for pedigree records.

The assignment of alleles to a breed origin was performed according to haplotypes defined by different core lengths. In a simulation study, Vandenplas et al. [5] assessed the impact of core length and observed higher assignment yield for haplotypes of longer core lengths. While this appears to be supported in our results, a core- and tail- length of 200 SNPs was observed as the optimal length for maximum assignment yield. Similarly, the impact of genetic relationship on assignment yield is comparable to values reported in simulated and empirical studies. Using simulated data, Vandenplas et al. [5], showed that a greater distance between breeds favourably affected the percentage of allele assigned, which is consistent with the highest percentage of allele assignment yield observed in crossbred5 (97%, Table 3) that are relatively distant to the local pure breeds.

The accuracy of allele assignment, both in the core-based (0.99, Table 3) and consensus-based (1.00, Table 4), across all scenarios was very high. This allele assignment accuracy is better than the results obtained from simulated (0.98) and empirical (88.57- 92.45) data [9]. The performance of the current algorithm is better than reported allele assignment accuracies of 96% using STRUCTURE 2.2 and 85% using GENECLASS 2 reported by Negrini et al. [15]. The relative performance improvement could be attributed to the optimization process of developing the current allele assignment algorithm. For example, the breed origin of alleles in crossbred animals was determined after an allele assignment was evaluated for every core and haplotype library in different scenarios to reach a consensus assignment. The choice of threshold for best SNP match in haplotypes can also affect the algorithm’s assignment yield and accuracy. Instead of using fixed allele frequency and best SNP matches to assign a breed origin to alleles, the observed expected trade-offs between assignment yield and accuracy (Fig. 6) have been optimized. When the best SNP match counts in haplotypes are too low, there will be a high assignment yield but low accuracy and vice versa. In the current simulated genotype data, the best SNP match count threshold of 50-60% appeared optimal.

Despite some suggestions to use haplotype instead of allele to reduce the effects of incorrect allelic assignments [5], the current algorithm was able to assign a breed origin to alleles as accurate as the assignment of a breed origin to haplotypes. The developed algorithm can be used to determine a breed origin of alleles in genomic predictions with models where breed-specific effects are required [16,17]. The developed algorithm can also be used in modelling breeding programmes of admixed populations. Accurate breed identification, on both the level of the individual and of the individual genetic variant is critical to achieving sustainable breed improvement. In the current simulation study, we developed an algorithm, which assigns haplotypes in crossbred dairy cows to the haplotypes of likely constituent breeds, i.e. either to exotic or local breeds. With high accuracy of assigning the breed of origin to alleles, we may be able to introduce a resilient or adaptive haplotype into the crossbred cows. In livestock, we infer haplotypes from multigenerational pedigrees from which tracing of breed origin of alleles can be challenging. With the developed algorithm, alleles in crossbred animals could be accurately assigned a breed of origin without the need for a multigenerational pedigree.

It’s important to acknowledge that the African dairy cattle populations are characterized by extensive crossbreeding involving many breeds of Taurine and Indicine origin. This broad genetic diversity may challenge the accurate estimation of SNP effects despite the accurate assignment of breed origin of alleles. While the BOA method relies on the recent local ancestry for each SNP marker allele, it ignores deeper ancestry, which is important for estimating SNP marker effects across many breeds with different genomic histories. Furthermore, the BOA method does not take full advantage of linkage information (correlation between nearby SNP markers) and does not fully reflect the underlying genomic history of a study population [18]. Future studies developing algorithms and methods that consider the BOA and the genomic history of individuals and that would work for any level of crossbreeding and admixture in a population will be needed.

## Conclusions

The developed algorithm assigns a breed origin to alleles with an accuracy of 99% in admixed animals from a crossbreeding programme designed to mimic breeding programmes in the LMICs. The algorithm is straightforward in its application and does not require prior knowledge of pedigree and relationships between crossbred and purebred animals, making it relevant and applicable in breeding programmes practised in LMICs. However, it should be noted that the algorithm was developed and tested on simulated data. Further studies are required to test and apply the algorithm on real data.

## Supporting information

R Scripts to simulate the genotypes

R scripts to phase the simulated genotypes

R scripts to assign a breed of origin to alleles of crossbred cattle

R scripts for consensus allele assignment

## List of abbreviations

ADGG: African Dairy Genetic Gains
BOA: Breed Origin of Allele
CTLGH: Centre for Tropical Livestock Genetics and Health
LMICs: Low- and middle-income countries
MaCS: Markovian Coalescent Simulator
SNP: Single Nucleotide Polymorphism

## Declarations

### Ethics approval and consent to participate

Not applicable

### Consent for publication

Not applicable

### Availability of data and materials

The scripts for data simulation and algorithm development are available [See Additional file 2, Script S2], [See Additional file 3, Script S3] and [See Additional file 4, Script S4].

## Authors’ contributions

BW, RCG, IH and JMH designed the study. BW performed the analyses and drafted the manuscript. IH has substantively revised the manuscript, addressed all the comments from co-authors and submitted the manuscript. GG and JMH supervised the study and contributed to the manuscript. All authors read and approved the final manuscript.

## Acknowledgements

We would like to thank Andrew Whalen, Appolinaire Djikeng, Joram M. Mwacharo and Jaap B. Buntjer for their supports during the study. Their fruitful discussions and advice led to the completion of the study.

## Additional files

Additional file 1 Script S1

File format: .txt

Title: R Scripts to simulate the genotypes.

Description: The Additional file 1 describes the scripts to simulate genotypes of the crossbred cattle.

Additional file 2 Script S2

Title: R scripts to phase the simulated genotypes File Format: .txt

Description: Additional file 2 describes the scripts to phase the simulated genotypes.

Additional file 3 Script S3

Title: R scripts to assign a breed of origin to alleles of crossbred cattle File Format: .txt

Description: Additional file 3 describes the scripts to assign a breed of origin to alleles of crossbred cattle

Additional file 4 Script S4

Title: R scripts for consensus allele assignment File Format: .txt

Description: Additional file 4 describes the Scripts for consensus allele assignment.

## References

1. Leroy G, Baumung R, Boettcher P, Scherf B, Hoffmann I. Review: sustainability of crossbreeding in developing countries; definitely not like crossing a meadow. Animal. 2016;10:262–73.

2. Marshall K, Gibson JP, Mwai O, Mwacharo JM, Haile A, Getachew T, et al. Livestock genomics for developing countries - African examples in practice. Front Genet. 2019; 10:297.

3. VanRaden PM, Cooper TA. Genomic evaluations and breed composition for crossbred US dairy cattle. Interbull Bull. 2015; 49: 19–23.

4. Kuehn LA, Keele JW, Bennett GL, McDaneld TG, Smith TPL, Snelling WM, et al. Predicting breed composition using breed frequencies of 50,000 markers from the US Meat Animal Research Center 2,000 Bull Project. J Anim Sci. 2011; 89:1742–50.

5. Vandenplas J, Calus MPL, Sevillano CA,Windig JJ, Bastiaansen JWM. Assigning breed origin to alleles in crossbred animals. Genet Sel Evol. 2016; 48:61.

6. Sevillano CA, Vandenplas J, Bastiaansen JWM, Bergsma R, Calus MPL. Genomic evaluation for a three-way crossbreeding system considering breed-of-origin of alleles. Genet Sel Evol. 2017; 49:75.

7. Guillenea A, Lund MS, Evans R, Boerner V, Karaman E. A breed-of-origin of alleles model that includes crossbred data improves predictive ability for crossbred animals in a multi-breed population. Genet Sel Evol. 2023; 55:34.

8. Eiríksson JH, Karaman E, Su G, Christensen OF. Breed of origin of alleles and genomic predictions for crossbred dairy cows. Genet Sel Evol. 2021; 53:84.

9. Sevillano CA, Vandenplas J, Bastiaansen JWM, Calus MPL. Empirical determination of breed-of-origin of alleles in three-breed cross pigs. Genet Sel Evol. 2016; 48:55.

10. Gaynor RC, Gorjanc G, Hickey JM. AlphaSimR: an R package for breeding program simulations. G3. 2021; 11:2.

11. Chen GK, Marjoram P, Wall JD. Fast and flexible simulation of DNA sequence data. Genome Res. 2009; 19:136–42.

12. R: a language and environment for statistical computing. Vienna: R Foundation for Statistical Computing; 2017.

13. Hickey JM, Kinghorn BP, Tier B, Wilson JF, Dunstan N, van der Werf JHJ. A combined long-range phasing and long haplotype imputation method to impute phase for SNP genotypes. Genet Sel Evol. 2011; 43:12.

14. Brown A, Ojango J, Gibson J, Coffey M, Okeyo M, Mrode R. Short communication: genomic selection in a crossbred cattle population using data from the dairy genetics East Africa project. J Dairy Sci. 2016; 99:7308–12.

15. Negrini R, Nicoloso L, Crepaldi P, Milanesi E, Colli L, Chegdani F, et al. Assessing SNP markers for assigning individuals to cattle populations. Anim Genet. 2009; 40:18–26.

16. Christensen OF, Madsen P, Nielsen B, Su GS. Genomic evaluation of both purebred and crossbred performances. Genet Sel Evol. 2014; 46:23.

17. Duenk P, Calus MPL, Wientjes YCJ, Breen VP, Henshall JM, Hawken R, et al. Estimating the purebred-crossbred genetic correlation of body weight in broiler chickens with pedigree or genomic relationships. Genet Sel Evol. 2019; 51:6.

18. Fan C, Mancuso N, Chiang CWK. A genealogical estimate of genetic relationships. Am J Hum Genet. 2022; 109: 812–24.

