## Supplementary material for "Accurate determination of breed origin of alleles in a simulated smallholder crossbred dairy cattle population": R Scripts to simulate the genotypes

#Scripts used to simulate the genotype and haplotype for the cattle breed 

#MaCS
#The split occurrences are given as a function of number of generations and effective population size (Ne).
#Split occurs = number of generations /4Ne
#For example, in the current study the value for the first split between the indigenous and exotic is 2.5, which assumes that the split occurred 1000 generations ago, with an effective population size of 100. 
#The split among the four indigenous breeds occurred 100 generations ago with an equal effective population size of 100. The detail for the MaCS parameters and the population demographic history is provided in the following R scripts used to simulate the genotype and haplotype data for an ancient cattle breed:

#Packages and R scrpits used to simulate the haplotypes of the local and exotic breeds
library(AlphaSimR)
macsCommand = function(size) {
  paste(1.00E+08,"-t",9.00E-06,"-r",3.60E-06,"-I 5",paste(rep(size/5,5),collapse=" "),
        "-eN 0.036 3.89 -eN 0.053 11.11 -eN 0.069 16.67 -eN 0.431 22.22 -eN 1.264 27.78 -eN 1.819 38.89 -eN 4.875 77.78 -eN 6.542 111.11 -eN 9.319 188.89 -eN 92.097 688.89 -ej 0.125 4 3 -ej 0.25 3 2 -ej 0.375 2 1 -ej 2.5 5 1 -s",sample.int(1e8,1))
  }

#Scrpit for simulation of 2 breeds, and terminal crossbreeding
genoError = 0; nSnp = 1000;  nInd = 2500; nChr = 1 #29; nGenerations=5;
nSnpPerChr = rep(nSnp, nChr) #Must be larger than nQtl

#simulation of founder population
founderPop = runMacs(nInd=nInd, nChr=nChr, segSites=nSnp,  nThreads = 4, manualCommand=macsCommand(2*nInd), manualGenLen=1)

#Only use animals from first breed for scaling traits
SP = SimParam$new(founderPop[1:500])
#track allele origin
SP$setTrackPed(TRUE) ; SP$setTrackRec(TRUE); SP$setGender(gender = "yes_sys")

#Set a SNP chip (not actually used in this simulation, just here for future reference)
SP$addSnpChip(nSnpPerChr = nSnp)
#The trait under selection has a small amount of dominance, which will create heterosis in crossbred animals
SP$addTraitAD(nQtlPerChr = nSnp,     mean = 0,   var = 1,   meanDD = 0.1, varDD = 0.1)
SP$setVarE(h2=0.3)

#initial breeds
A = newPop(founderPop[1:500])
B = newPop(founderPop[501:1000])
C = newPop(founderPop[1001:1500])
D = newPop(founderPop[1501:2000])
exotic = newPop(founderPop[2001:2500])

localBreeds = vector("list", nGenerations)
exoticBreeds=vector("list", nGenerations)
crossbred = vector("list", nGenerations)

localBreeds[[1]] = c(A, B, C, D)
exoticBreeds[[1]] =  exotic

#random mating of within local and exotic animals
for(i in 2:nGenerations){
localBreeds[[i]] =  randCross(localBreeds[[i-1]],nCrosses=2000)
exoticBreeds[[i]] = randCross(exoticBreeds[[i-1]],nCrosses=2000)
}

#select local females from last generation and exotic males
exoticBulls = selectInd(exotic, nInd=25, gender = "M")
localCows= selectInd(localBreeds[[nGenerations]], nInd=1000,gender = "F")

#produce first crossbred of local females and exotic males
crossbred[[1]] = randCross2(females=localCows, males=exoticBulls, nCrosses=2000)
#produce crossbred of crossbredfemale and exotic males in each generation for upto 5 generations (round of selctions)
for(i in 2:nGenerations) {
  crossbred[[i]] = randCross2(females=selectInd(crossbred[[i-1]], nInd=1000, gender = "F"), males=selectInd(exoticBreeds[[i-1]], nInd=25, gender="M"), nCrosses=2000)
}

write.table2 = function(x, file, ...) {
  write.table(x, file, col.names=F, row.names=F,quote=F,...)
}

for(i in 1:nGenerations){
  hapcrossbred = pullIbdHaplo(crossbred[[i]])
  write.table2(hapcrossbred, paste0("hapcrossbred", i))
}
hapexotic = pullIbdHaplo(exoticBreeds[[i]])
haplocal = pullIbdHaplo(localBreeds[[i]])

exoticout=exoticBreeds[[nGenerations]]
localout=localBreeds[[nGenerations]]

writePopulation = function(pop, prefix) {
	pedigree = data.frame(pop@id, pop@father, pop@mother, stringsAsFactors = FALSE)
	aa = AlphaSimR::pullSnpGeno(pop, 1)
	# bb = AlphaSimR::pullSnpHaplo(pop, 1)
	write.table2(pedigree, paste0("pedigree.", prefix))
	write.table2(cbind(pedigree[,1], aa), paste0("genotype.", prefix))
}

writePopulation(localout, "localbred")
writePopulation(exoticout, "exoticbred")

for(i in 1:nGenerations) {
  writePopulation(crossbred[[i]], paste0("crossbred", i))
}


# References:
# Gaynor, R., G. Gorjanc, D. Wilson, D. Money & J. Hickey (2019) AlphaSimR: Breeding Programme Simulations. URL https://CRAN. R-project. org/package= AlphaSimR, r package version 0.9. 0.
# Wang, J., Santiago, E. & Caballero, A. Prediction and estimation of effective population size.ÊHeredityÊ117, 193Ð206 (2016). https://doi.org/10.1038/hdy.2016.43
